# PINK1 and parkin shape the organism-wide distribution of a deleterious mitochondrial genome

**DOI:** 10.1101/576165

**Authors:** Arnaud Ahier, Nadia Cummins, Chuan-Yang Dai, Jürgen Götz, Steven Zuryn

## Abstract

In multiple species, certain tissue types are prone to acquiring greater loads of mitochondrial genome (mtDNA) mutations relative to others, however the mechanisms that drive these heteroplasmy differences are unknown. We found that the conserved PTEN-induced putative kinase (PINK1/PINK-1) and the E3 ubiquitin-protein ligase parkin (PDR-1), which are required for mitochondrial autophagy (mitophagy), underlie stereotyped differences in heteroplasmy of a deleterious mitochondrial genome mutation (ΔmtDNA) between major somatic tissues types in *Caenorhabditis elegans*. We demonstrate that tissues prone to accumulating ΔmtDNA have lower mitophagy responses than those with low mutation levels, such as neurons. Moreover, we show that ΔmtDNA heteroplasmy increases when proteotoxic species that are associated with neurodegenerative disease and mitophagy inhibition are overexpressed in the nervous system. Together, these results suggest that PINK1 and parkin drive organism-wide patterns of heteroplasmy and provide evidence of a causal link between proteotoxicity, mitophagy, and mtDNA mutation levels in neurons.

## Introduction

Mitochondria act as central hubs for cellular bioenergetics, macromolecule precursor synthesis, redox balance, Ca^2+^ handling, apoptosis, and immunity [1, 2]. They house their own genome (mtDNA), as well as RNA and protein-synthesizing systems, which together code and coordinate the assembly of core subunits of oxidative phosphorylation (OXPHOS). Mutations in the mtDNA that perturb the assembly of OXPHOS enzymes can cause devastating metabolic disorders [3]. The level of heteroplasmy of a pathogenic mtDNA mutation correlates with the severity of the clinical phenotype and mosaic distributions of mutations across an individual have been reported to lead to organ-specific dysfunction [4]. Heteroplasmy and mosaicism are therefore important determinants of mitochondrial disease pathophysiology.

The composition of the mitochondrial DNA (mtDNA) in each cell is in constant flux through processes of mutation, replication, and degradation. Mitotic segregation of mitochondria may result in asymmetric proportions of heteroplasmic allelic variants in descendant cellular lineages [5]. Alternatively, due to the high rate and relaxed replication of mtDNA, mosaicism can arise through clonal expansion and subsequent genetic drift between adjacent cells and tissues over time [6]. Indeed, there is continuous replication of mtDNA in all tissues, allowing variations in heteroplasmy to even develop between post-mitotic cells. Although there is evidence of selectivity [7], it is generally assumed that mitotic segregation and genetic drift are largely stochastic processes and therefore lead to random variation in mtDNA heteroplasmy and mosaicism within individuals [4]. Evidence in several species, however, suggests that differences in heteroplasmy between certain cell types is not random and may instead be deterministic. For example, relative to the rest of the human brain, the caudate, putamen and substantia nigra regions show an age-dependent increase in the heteroplasmy levels of the mtDNA^4977^ deletion (a mutant genome harbouring a 4,977bp deletion) [8]. Heteroplasmy differences between tissues have also been observed in *C. elegans*. Here, mtDNA molecules harbouring a 3.1 kb deletion (ΔmtDNA) accumulate at different heteroplasmy levels in distinct somatic and germ lineages over time [9, 10]. How different cell, tissue, or organ types reproducibly develop higher heteroplasmy levels than others is unclear.

By purifying and analysing mitochondria from different cell types, we show that PINK-1 and PDR-1 are required to set heteroplasmy differences between major somatic tissue groups in *C. elegans* and that removal of these genes equalises ΔmtDNA heteroplasmy across the organism. Mechanistically, our evidence indicates that heteroplasmy is lowered by PINK-1 and PDR-1 through the selective removal of ΔmtDNA, but to varying degrees of efficiency in different cell types with neurons, intestinal cells, and epidermal cells being more amenable to ΔmtDNA clearance than body wall muscle cells. As a result, body wall muscle cells accumulate ΔmtDNA at higher levels than the other tissues. Interestingly, we find that neurons, which have the greatest dependency upon PINK-1 and PDR-1 for maintaining low ΔmtDNA heteroplasmy levels, are susceptible to increases in heteroplasmy caused by overexpressing either tau or polyglutamate repeats in the nervous system, proteotoxic species associated with neurodegenerative disease and mitophagy inhibition. Together, our results uncover a nuclear-encoded mechanism that determines organism-wide patterns of mtDNA heteroplasmy and establishes links between proteotoxicity, mitophagy, and mtDNA mutation levels in neurons.

## Results

To understand the mechanisms that underlie the divergence in heteroplasmy levels between different tissue types within individuals, we took advantage of a strain of *C. elegans* that harbours a 3.1 kb deletion in its mitochondrial genome (*uaDf5*, also called ΔmtDNA). This mutation exists in stable heteroplasmy of 50-60% (Figure S1A) and is deleterious for mitochondrial function as it disrupts four mtDNA genes that encode core subunits of the respiratory chain (Figure S1B). Worms harbouring the *uaDf5* allele at around 60% heteroplasmy display reductions in basal oxygen consumption and total respiratory capacity as well as constitutive induction of the mitochondrial unfolded protein response (UPR^mt^) [11, 12]. Because cellular heteroplasmy is determined by the net result of the propagation and degradation of mtDNA mutations, we investigated whether mitophagy, a process that can selectively destroy depolarized mitochondria harbouring mtDNA mutations, such as ΔmtDNA [11–13], may underlie tissue-specific heteroplasmy levels. To test this idea, we introduced the ΔmtDNA mutation into double-mutant strains with null deletions in the genes *pink-1* and *pdr-1* [14], key mediators of mitophagy in multiple species including *C. elegans* [15–18]. PINK1/PINK-1 can induce mitophagy by accumulating on the outer mitochondrial membrane of defective organelles, where it recruits the E3 ubiquitin ligase parkin/PDR-1 [19]. Parkin/PDR-1 decorates the surface of these mitochondria with polyubiquitin chains, thereby inducing engulfment by autophagosomes and subsequent destruction following fusion with lysosomes [20]. In some invertebrate species such as *Drosophila*, parkin can act independently of PINK1 during mitophagy [21], prompting us to use *pink-1(tm1779);pdr-1(gk448)* double mutants to unambiguously disrupt mitophagy in *C. elegans. Pink-1(tm1779);pdr-1(gk448)* double mutants harbouring heteroplasmic ΔmtDNA were crossed to a series of transgenic strains carrying a *TOMM-20::mKate2::HA* transgene under the control of either the promoter of the gene *myo-3, ges-1, rgef-1*, or *dpy-7* which selectively drive body wall muscle, intestinal, panneuronal, and epidermal expression, respectively (Figure 1A). We also generated a similar strain which carried the transgene *TOMM-20::mKate2::HA* under the ubiquitous control of the promoter of *eft-3*. Intact mitochondria were isolated from each of the strains above, as well as wild-type animals in which mitophagy was unperturbed. This was performed by using immunoprecipitation against the HA epitope, which labelled the outer surface of the mitochondria because of its attachment to the C-terminus of TOMM-20 [9]. In doing so, we were able to selectively purify mitochondria from each tissue type in which the *TOMM-20::mKate2::HA* transgene was expressed.

**Figure 1.**
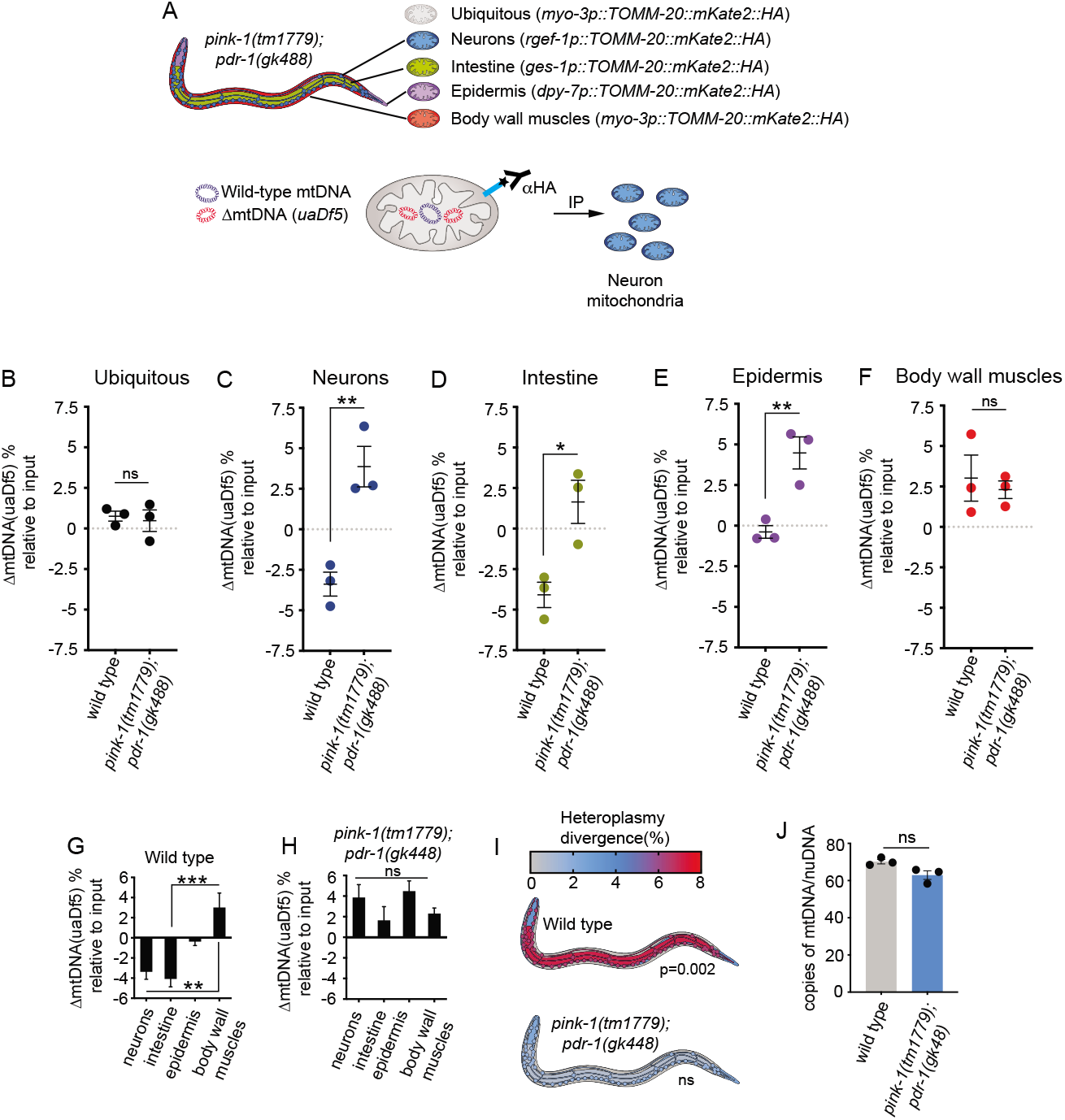
*Pink-1* and *pdr-1* are required for establishing differences in ΔmtDNA heteroplasmy between major tissue types. (**A**) Overview of mitochondrial immunoprecipitation from specific tissues in *pink-1(tm1779);pdr-1(gk488)* double mutants harbouring ΔmtDNA. (**B-F**) Relative heteroplasmy differences in mitochondria isolated from each cell type in wild type and *pink-1(tm1779);pdr-1(gk488)* backgrounds. Bars are means ± s.e.m. of three independent biological replicates (dots) consisting of ~10,000 animals each. * = *p*< 0.05; ** = *p*< 0.01. *P* values were determined using one-way ANOVA and Tukey’s correction. Comparison of heteroplasmy between tissues in (**G**) wild-type animals and (**H**) *pink-1(tm1779);pdr-1(gk488)* double mutants indicates that inter-tissue differences in ΔmtDNA heteroplasmy is abolished by removing *pink1* and *pdr-1*. (**I**) Graphical representation of heteroplasmy divergence (relative to body wall muscle) between tissues in each background. The significance of heteroplasmy variation between tissues was calculated using one-way ANOVA. ** = *p*< 0.01; *** = *p*< 0.001. *P* values were determined using one-way ANOVA and Tukey’s correction. (**J**) Quantitative PCR analysis of mtDNA copy number in L4 wild type and *pink-1(tm1779);pdr-1(gk488)* double mutants. Columns are means ± s.e.m. of three independent biological replicates (dots), each consisting of 10 animals, analysed with Student’s t-test.

Heteroplasmy comparisons in animals expressing *eft-3p::TOMM-20::mKate2::HA* revealed that there was no relative difference in ΔmtDNA levels between immunoprecipitated mitochondria and total (non-affinity purified input sample) mitochondria (Figure 1B). This was as expected and indicated that the mitochondrial isolation procedure used did not artificially alter heteroplasmy level measurements made in either wild-type or *pink-1(tm1779);pdr-1(gk448)* backgrounds. Comparisons in relative ΔmtDNA levels made between mitochondria purified from each tissue type in wild-type worms revealed that neurons, intestinal cells, epidermal cells, and body wall muscle cells displayed distinct and reproducible differences in ΔmtDNA heteroplasmy relative to each other (Figures 1C-F), with muscle cells always harbouring the highest load of mutations of any somatic tissue type studied (Figure 1G)[9]. Interestingly, these intertissue differences in heteroplasmy were abolished in *pink-1(tm1779);pdr-1(gk448)* mutants (Figure 1H and 1I), suggesting that the activities of *pink-1* and *pdr-1*, two nuclear-encoded genes, were required for setting distinct and stereotyped patterns of heteroplasmy in major somatic tissues of *C. elegans*. This equalisation effect wasn’t caused by global changes in mtDNA copy number, as *pink-1(tm1779);pdr-1(gk48)* mutants had normal levels of mtDNA (Figure 1J). At the level of individual tissues in *pink-1(tm1779);pdr-1(gk488)* double mutants, we observed increases in heteroplasmy in neurons, intestinal cells, and epidermal cells (Figure 1C-E), but no alteration in ΔmtDNA levels in body wall muscle cells (Figure 1F). This suggested that *pink-1* and *pdr-1* acted only in select cell types to remove ΔmtDNA, leading to stereotyped differences in heteroplasmy between tissues over time.

Because these results indicated that *pink-1* and *pdr-1* did not influence heteroplasmy in muscle cells, we hypothesised that the genes were either inactive or that they had different functions that did not include ΔmtDNA clearance in this tissue type. However, we found that *pink-1* was expressed in body wall muscle cells, as indicated by a *pink-1p::PINK-1::GFP* [22] transgene in live animals (Figure 2A). Because a translational GFP::PDR-1 reporter is also expressed in body wall muscles in *C. elegans* [14], it is likely that PINK-1-PDR-1-mediated mitophagy is active in this tissue. In *C. elegans*, mitophagy can be monitored through the colocalization of mitochondria with autophagosomes, which defines one of the terminal stages of mitophagy prior to lysomal fusion and the destruction of encapsulated mitochondria [16, 23]. By quantifying the colocalization of *myo-3p*::TOMM-20::mKate2::HA and *lgg-1p:*:LGG-1::GFP [24], which label muscle mitochondria and autophagosomes, respectively, we determined that mitophagy was inhibited in the body wall muscle cells of *pink-1(tm1779);pdr-1(gk488)* worms (Figure 2B). Indeed, in these mutants, using multiple fluorescent markers (*myo-3p::TOMM-20::mKate2::HA* and *myo-3p::^MTS^RFP*) and three-dimensional reconstruction of high magnification images, we also observed a dramatic increase in total mitochondrial muscle volume (Figure 2C, D and Figure S3), suggesting that the mitochondrial network increased in size in the absence of mitophagy-mediated turnover. Together, these results indicate that *pink-1* and *pdr-1* are active and required for mitophagy in muscle cells in *C. elegans*, consistent with previous findings [16].

**Figure 2.**
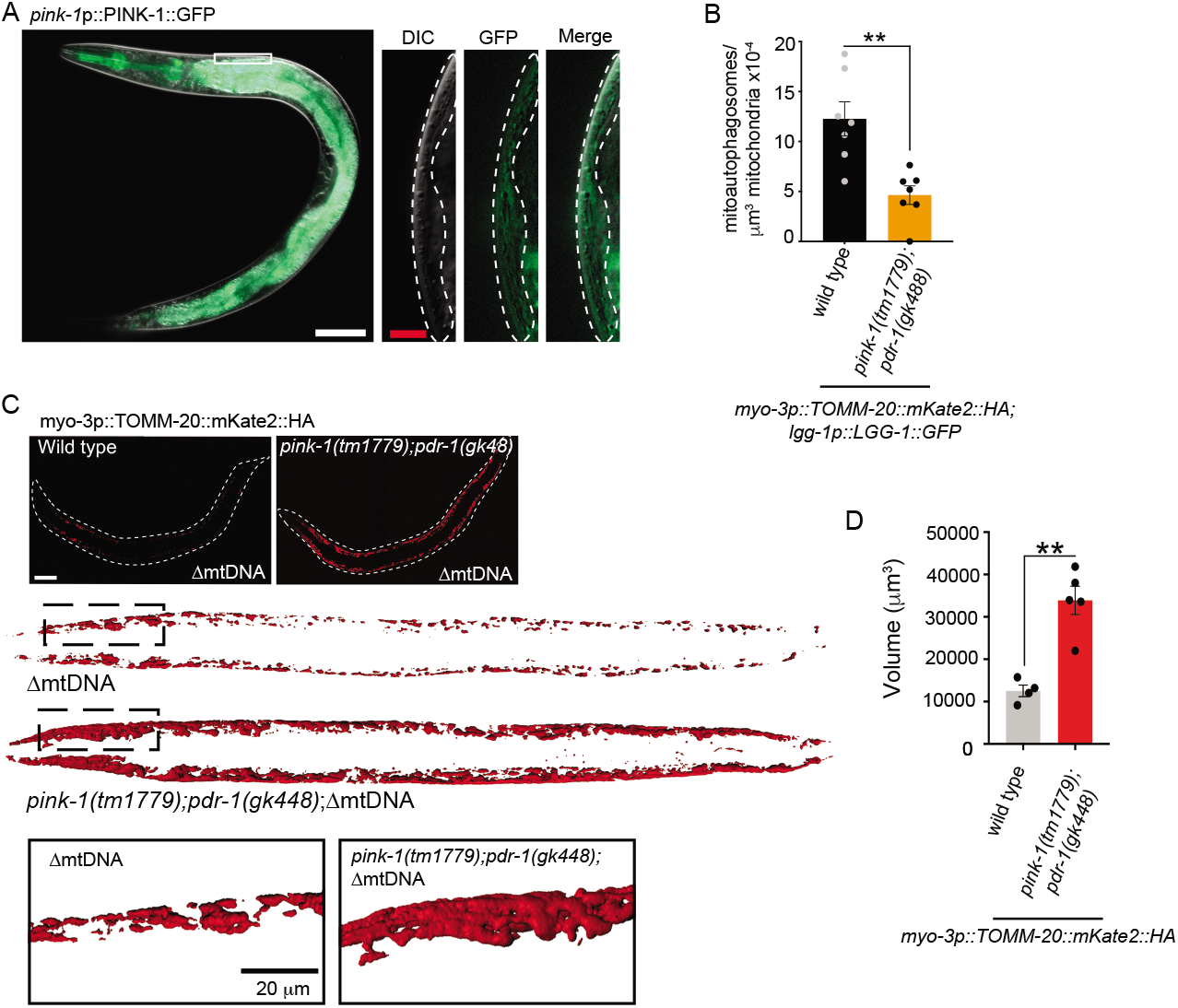
*Pink-1* and *pdr-1* are required for mitophagy in body wall muscle cells. **(A)** PINK-1 translational reporter is expressed in most somatic tissues in *C. elegans*, including the body wall muscle. White box is magnified in the panels on the left and body muscle cells are outlined in white dashed lines. White scale bar, 50μm; red scale bar, 5μm. (**B**) Quantification of mitophagy in body wall muscle cells. Columns are means ± s.e.m. of independent animals (grey and black dots). ** = *p*< 0.01. *P* values were determined using Student’s t-test. (**C**) Representative images and three-dimensional reconstructions of the *C. elegans* body wall muscle mitochondrial network. These representative figures have been constructed from high magnification (40x) z-stack images of live animals stitched together and three-dimensionally reconstructed. (**D**) Quantification of body wall muscle mitochondrial volume. Columns are means ± s.e.m. of individual animals (black dots). ** = *p*< 0.01, *P* values were determined using Student’s t-test.

Mitophagy is initiated by the stabilisation of PINK1 on the surface of depolarised mitochondria. However, although mitochondria are ubiquitous, they can exhibit cell-type specific morphologies, dynamics, and functions and as such, mitochondria from distinct cell-types may respond physiologically differently to the presence of ΔmtDNA in a manner that impinges on mitophagy and therefore ΔmtDNA clearance. For example, if the mitochondrial membrane potential (MMP) is unaffected by ΔmtDNA in muscle cells, it is possible that mitochondria harbouring ΔmtDNA may not be recognised and removed via mitophagy. To investigate this possibility further, we compared the effects of ΔmtDNA in body wall muscle cells and neurons, two cell types with opposite heteroplasmy trends (Figure 1C and F). Although gross mitochondrial morphology and volume appeared to be unaffected by ΔmtDNA (Figure S3), we found that ΔmtDNA induced a decrease in the MMP in both muscle and neural mitochondria to the same degree (Figure 3A and B). Furthermore, we found that total mtDNA copy number increased in mitochondria purified from muscle cells and neurons, which is likely to be a compensatory response to mitochondrial dysfunction caused by the presence of ΔmtDNA (Figure 3C). This suggests that ΔmtDNA invokes comparable levels of mitochondrial dysfunction and depolarisation in muscle cells and neurons and that tissue-specific differences in heteroplasmy, as determined by *pink-1* and *pdr-1* activities, are not attributable to intrinsic differences in mitochondrial physiology between cell types.

**Figure 3.**
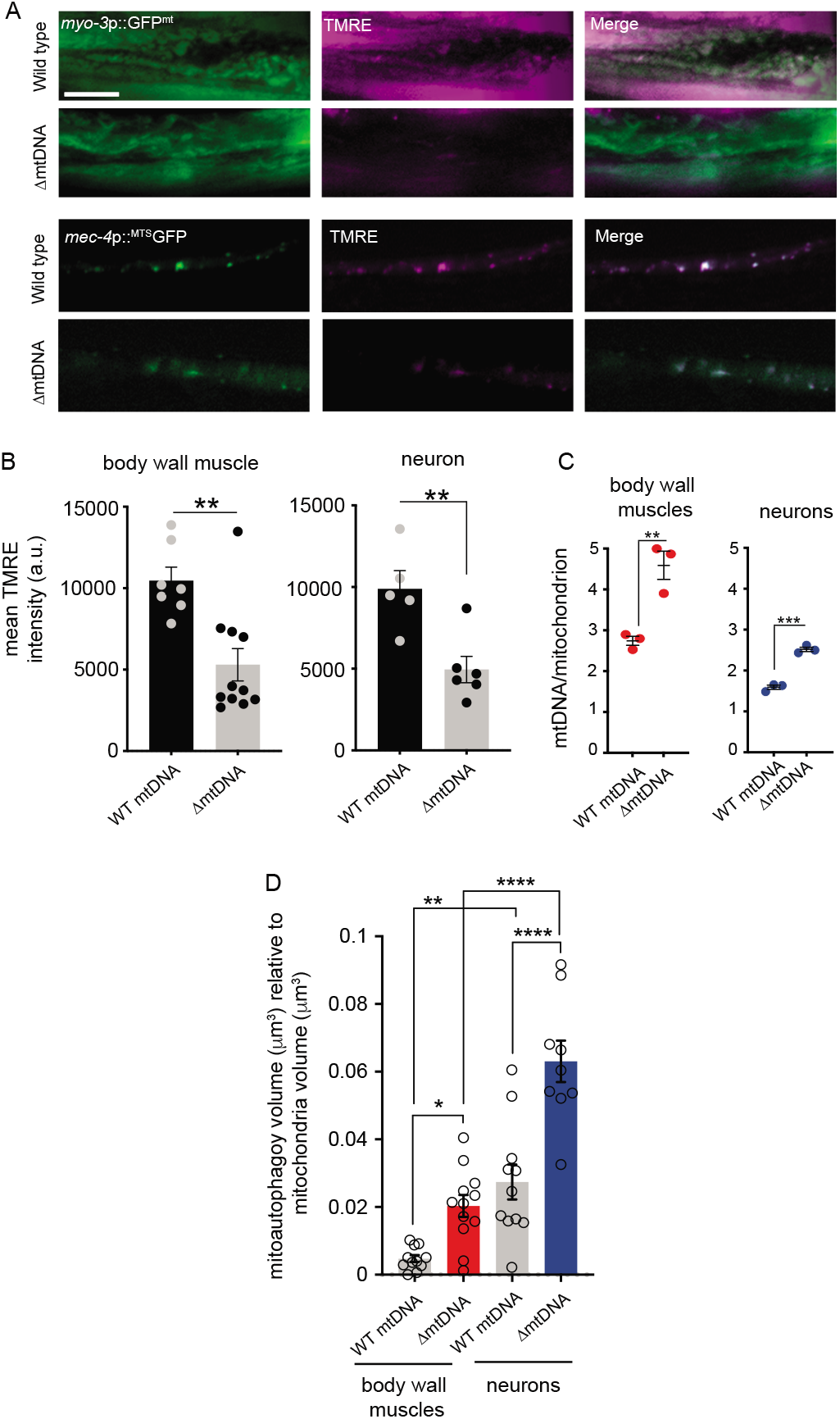
ΔmtDNA induces mitochondrial dysfunction and mitophagy in muscle cells and neurons. (**A**) Representative micrographs taken of live animals expressing tissue-specific GFP reporters (*myo-3*p::GFP^mt^, body wall muscle and *mec-4*p::^MTS^GFP, mechanosensory neurons) localised to the mitochondria and stained with the mitochondrial membrane potential sensitive dye, TMRE (pseudo-coloured magenta). Scale bar, 10μm. (**B**) Quantification of mean TMRE intensity from mitochondria of each tissue type. a.u., arbitrary units. Columns are means ± s.e.m., each dot is the mean TMRE intensity calculated from an independent animal. ** = *p*< 0.01. *P* values were determined using Student’s t-test. (**C**) Quantification of mtDNA molecules per mitochondrion. Each dot represents an independent experiment in which the average number of mtDNA molecules per mitochondrion (n > 1,000 mitochondria per sample) were calculated. Bars are means of three biologically independent experiments ± s.e.m. ** = *p*< 0.01; *** = *p*< 0.001. *P* values were determined using one-way ANOVA and Tukey’s correction. (**D**) Quantification of mitophagy in body wall muscle cells and neurons. Columns are means ± s.e.m., each dot represents an independent animal analysed. * = *p*< 0.05; ** = *p*< 0.01; *** = *p*< 0.001; **** = *p*< 0.0001. *P* values were determined using one-way ANOVA and Tukey’s correction.

Although *pink-1* and *pdr-1* are expressed and are required for mitophagy in both body wall muscle (Figure 2B) and neurons, as we showed previously in *C. elegans* [15], it is possible that mitophagy responds differently to the presence of ΔmtDNA in each cell type, which may affect its clearance. We compared tissue-specific levels of mitophagy in the nervous system and body wall muscle in wild-type animals and animals harbouring ΔmtDNA by determining the percentage overlap (volume:volume) of mitochondria (muscle, *myo-3p::TOMM-20::mKate2::HA*; neural *rgef-1p::TOMM-20::mKate2::HA*) to autophagosomes (*lgg-1p::LGG-1::GFP*). Interesting, we found that mitophagy was induced by the presence of ΔmtDNA in both cell types, suggesting that mtDNA mutations can directly activate mitochondrial clearance pathways *in vivo* (Figure 3D). More importantly, we found that relative to mitochondrial volume, there was much more basal mitophagy in neurons compared to muscle cells. Moreover, although both tissue types responded to the presence of ΔmtDNA by increasing mitophagy, the neural response was much greater than that in muscle (Figure 3D). This suggests that mitophagy has a greater capacity to respond to ΔmtDNA in the nervous system than in muscle and could therefore explain why neurons have lower heteroplasmy levels than muscle and why mutations in *pink-1* and *pdr-1(gk488)* have a greater effect on heteroplasmy in the nervous system.

Because our results indicated that ΔmtDNA strongly induces mitophagy in the neurons and that the activity of PINK-1-PDR-1-mediated mitophagy is important in the nervous system for maintaining low levels of heteroplasmy, we next determined whether other cellular conditions that perturbed mitophagy could raise ΔmtDNA levels in these cells types. We recently demonstrated that proteotoxic tau, a protein that forms aggregates in tauopathies such as Alzheimer’s disease [25], perturbs parkin translocation to the mitochondria, and that human tau expressed in the *C. elegans* nervous system inhibits mitophagy [15]. To determine whether this increased heteroplasmy levels in neurons, we constructed strains harbouring ΔmtDNA and expressing human tau (hTau) in the nervous system (*aex-3p::hTau*) [15] as well as the neuron-specific *rgef-1p::TOMM-20::mKate2::HA* transgene (Figure 4A). We then immunoprecipitated neuronal mitochondria in these strains and found that hTau significantly increased heteroplasmy (Figure 4B), suggesting that the expression of mitophagy-inhibiting proteins can increase ΔmtDNA levels in neurons. Other proteotoxic species have also recently been found to inhibit mitophagy in neurons. The expanded polyglutamine (polyQ) tract in mutant forms of the Huntingtin protein (Htt) that cause Huntington disease were recently reported to impede mitophagy in differentiated striatal neurons [26]. Indeed, we found that the expression of a polyglutamine (polyQ) tract (*rgef-1p::Q40::YFP*) [27] in the *C. elegans* nervous system recapitulated the effect of hTau and increased ΔmtDNA heteroplasmy in neurons (Figure 4B). These results suggest that diverse proteotoxic threats associated with a broad range of neurodegenerative diseases may promote the accumulation of mtDNA deletions in the nervous system.

**Figure 4.**
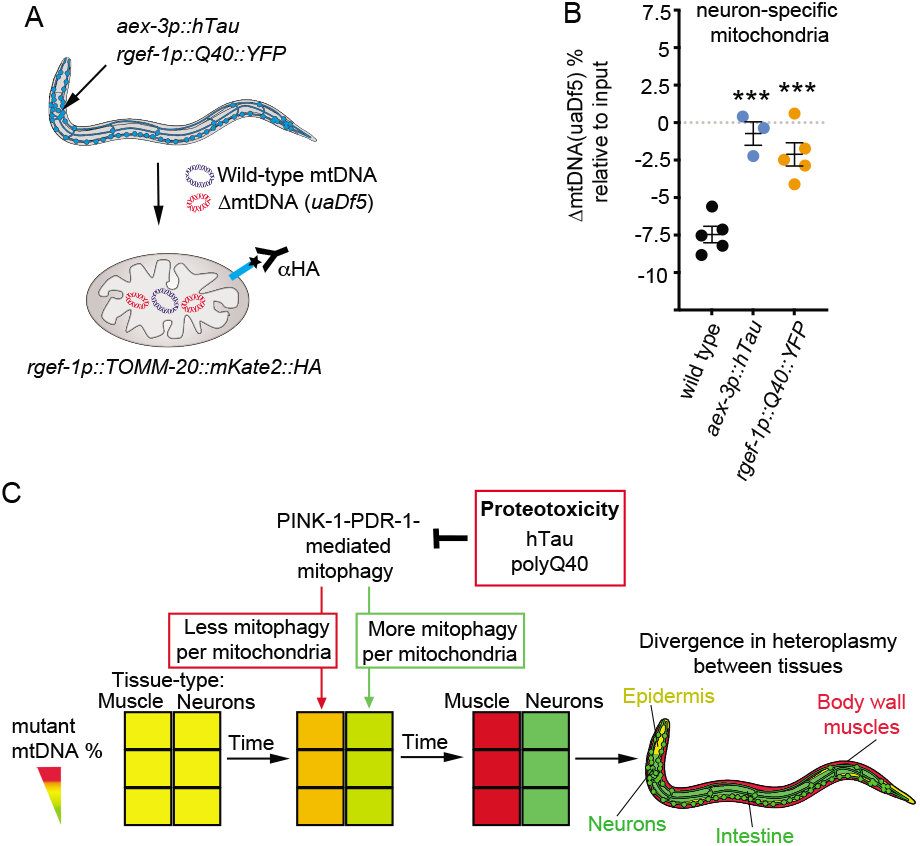
Proteotoxicity increases ΔmtDNA heteroplasmy in neurons. (**A**) Overview of the genetic strains used to isolate mitochondria from neurons overexpressing proteotoxic species. (**B**) Expression of either human tau (hTau) or a polyglutamine tract (Q40) in neurons increases ΔmtDNA levels in the nervous system. Bars are means ± s.e.m. of at least three independent biological replicates (dots). *** = *p*< 0.001. *P* values were determined using one-way ANOVA and Tukey’s correction. (**C**) Proposed model.

## Discussion

Stochastic processes such as mitotic segregation and genetic drift influence cellular heteroplasmy [4]. However, certain tissues are prone to accumulating mtDNA mutations, implying that cell-type specific events deterministically set heteroplasmy levels. Our results suggest that cell-type differences in the activity and responsiveness of mitophagy, mediated through *pink-1* and *pdr-1*, contribute to the intertissue divergence in heteroplasmy of a deleterious mitochondrial genome. We showed that deletion of *pink-1* and *pdr-1* equalised heteroplasmy levels across tissues that normally display stereotypical differences in ΔmtDNA heteroplasmy. Comparisons between body wall muscle cells and neurons revealed that although active in both tissues, mitophagy responded to ΔmtDNA at greater levels relative to total mitochondrial volume in neurons than muscle, leading to higher ΔmtDNA heteroplasmy in muscle. Furthermore, we have demonstrated that proteotoxic threats that can perturb mitophagy in neurons can increase heteroplasmy levels in the nervous system. Taken together, our results suggest a model (Figure 4C) whereby organism-wide patterns of heteroplasmy of mtDNA deletions are determined by the activity of PINK-1-PDR-1-mediated mitophagy and that genetic perturbations and disease-related proteotoxic threats that perturb mitophagy can increase the abundance of mtDNA mutations in the nervous system of an organism.

In humans, large mtDNA deletions have been shown to accumulate in specific regions of the brain during ageing [8]. Although there is strong evidence that PINK1-parkin-mediated mitophagy occurs in mammalian neurons [28, 29], it is not known whether differences in the activity of mitophagy plays a role in specifying the heteroplasmy levels of mtDNA deletions within distinct regions of the brain. In mice homozygous for a proofreading deficiency in DNA polymerase γ, which induces a high level of mtDNA single nucleotide mutations [30, 31], parkin knockout significantly increases the levels of pathogenic mtDNA mutations in the striatum [32]. Moreover, mutations in Pink1 [33] and parkin [34] can lead to early onset Parkinson’s disease, and substantia nigra neurons in Parkinson’s disease patients have high levels of mtDNA deletions [35, 36]. Although we did not examine subsets of neurons in this study, our results suggest that neurons in general are the most vulnerable to increases in heteroplasmy when mitophagy is perturbed. Therefore, our work demonstrates that mutations in Pink1 and parkin have a widely conserved effect across phyla of enhancing the accumulation of mtDNA deletions in neurons, which may perturb their function and cause disease. It will be interesting to determine whether these principles are specific to mtDNA deletions or other types of mtDNA mutations as well.

In addition to Pink1 and parkin mutations, we found that toxic proteins associated with neurodegenerative diseases other than Parkinson’s disease also increase heteroplasmy. Overexpression in neurons of hTau, as well as a polyQ tract, significantly increased ΔmtDNA heteroplasmy levels in the *C. elegans* nervous system. To our knowledge, this represents the first reported incidence of proteotoxicity increasing the cellular load of mtDNA mutations. Interestingly, similar proteotoxic species have been recently reported to inhibit mitophagy in neurons. Via its projection domain, hTau can associate with Parkin in the cytoplasm and specifically impair its translocation to the surface of defective mitochondria in human neuroblastoma cells [15]. In addition, a polyQ tract within expanded mutant versions of the Huntingting protein can prevent the formation of the Beclin1-Vps34 initiation complex, negatively affecting the efficiency of mitophagy in post-mitotic striatal neurons [26]. These results demonstrate that perturbing mitophagy indirectly with proteotoxicity has the same effect as perturbing mitophagy genetically through mutations in *pink-1* and *pdr-1*. They also demonstrate that toxic proteins associated with diverse neurodegenerative diseases can modify the genetic landscape of the mitochondrial genome in neurons and provide evidence that two of the most common hallmarks of neurodegeneration - proteotoxicity and mitochondrial dysfunction - may converge at the level of the mitochondrial genome.

## Materials and Methods

### *C. elegans* strains and culture

LB138 *him-8(e1489) IV; uaDf5/+*; EG8078 *oxTi185 I; unc-119(ed3) III*; BR4006 *pink-1(tm1779) II; byEx655 [pink-1p::pink-1::GFP; myo-2p::mCherry; herring sperm DNA]*; VC1024 *pdr-1(gk448) III*; AM101 *rmIs110 [rgef-1p::Q40::YFP]*, SJ4103 *zcIs14 [myo-3p::GFP^mt^]*, MAH235 *sqIs13[lgg-1p::gfp::lgg-1 + odr-1p::RFP]* [24] and wild-type (N2) were acquired from the *Caenorhabditis* Genetics Center (CGC), University of Minnesota, USA, which is funded by the NIH Office of Research Infrastructure Programs (P40 OD010440). APD178 *apdIs11 [aex-3p::tau4RIN + coel::gfp]* were kindly provided by Dr H. Nicholas (The University of Sydney, Sydney, Australia). QH5425 *vdIs30 [unc-25p::MTS::mRFP + odr-1p::dsRed]* was kindly provided as a gift by Dr M. Hilliard and J. Chaplin (The University of Queensland, Queensland Brain Institute, Brisbane, Australia). *C. elegans* culture and maintenance were performed according to standard protocols [37]. The transgenic strains used for cell-specific mitochondrial purification experiments, *foxSi16[myo-3p::TOMM-20::mKate2::HA], foxSi37 [ges-1p::TOMM-20::mKate2::HA], foxSi41[dpy-7p::TOMM-20::mKate2::HA], foxSi44[rgef-1p::TOMM-20::mKate2::HA], foxSi75[eft-3p::TOMM-20::mKate2::HA]* and *foxSi2[myo-3p:: ^MTS^RFP]* were used were generated as previously described [9].

### Cell-specific mitochondrial affinity purification (CS-MAP)

CS-MAP was performed as previously described [9]. Heteroplasmy in mitochondria derived from different cell types was investigated across very large populations of pooled animals (>10,000 individuals). This approach enabled population-scale trends to be resolved by overcoming confounding influences that can be introduced by the presence of stochastic inter-individual variations. Briefly, 10,000 to 20,000 L4 staged worms were grown on 2 x 150 mm NGM plates seeded with *E. coli* OP50 bacteria. We chose L4 animals as it represents a stage in which all somatic tissues are fully formed (e.g. nervous system and epidermal seam cells) and developing embryos are not yet present in the gonads of animals. Worms were homogenized into a hypotonic buffer [50 mM KCl, 110 mM mannitol, 70 mM sucrose, 0.1 mM EDTA (pH 8.0), 5 mM Tris-HCl (pH 7.4)] with a dounce homogenizer and the subsequent crude mitochondrial fraction was enriched by differential centrifugations. The mitochondria were then isolated from this fraction using anti-HA (influenza hemagglutinin) magnetic beads (Pierce). The beads were washed and resuspended in 20 μl of hypotonic buffer. All CS-MAP experiments were repeated in at least three independent experiments and data for each genetic background were obtained at similar times to those previously reported for control wild-type backgrounds [9]. To eliminate the possibility of any population variability affecting our analyses, we compared mitochondria purified from each tissue to the total mitochondria (homogenate) from the same samples.

### mtDNA extraction and quantitative PCR of heteroplasmy

For all mtDNA analyses, total input and affinity-purified mitochondria were lysed and extracted using phenol-chloroform-isoamyl alcohol, followed by an ethanol precipitation. Extracted DNA was resuspended in 10 mM Tris-HCl (pH 8), 0.1 mM EDTA. Multiplex quantitative PCR was then performed using the Luna Universal Probe qPCR Master Mix (ref M3004, NEB), a Roche LightCycler or Rotor Gene Q real-time PCR (Qiagen) machine and Rotor Gene Q pure detection software (V2.3.1). The primers 5’-cgtaagaaaatcaaaatatggtataattgg-3’, 5’-aaaatgttacgatccttattaataaagc-3’, 5’-gcagagatgtttattgaagctgac-3’ and the probes 5’-HEX/tgaggccag/ZEN/ttcatattgttccaga gtg/IABKFQ-3’ (Iowa Black FQ) and 5’-6-FAM/ccatccgtg/ZEN/ctagaagacaaagaatttc/IABKFQ-3’ were used to quantify the wild-type mtDNA and ΔmtDNA levels. Calculations of mtDNA copies per mitochondrion were derived from the quantification of purified organelles in individual samples, as previously performed [9]. Control wild-type values have been previously reported by Ahier et al., 2018 [9].

### Quantification of mtDNA copy number

Ten (10) L4 worms from each genetic background were lysed in 10 μl of standard lysis buffer supplemented with 200□ng□ml-1 proteinase K and incubated at 65°C for 2□h, followed by heat-inactivation at 95□°C for 15□min. A 1/50 dilution of the lysate was used for subsequent quantitative PCR. The Sensifast SYBR no-rox Kit (ref. BIO-98005, Bioline) and a Rotor Gene Q Real-Time PCR (Qiagen) Machine, with Rotor Gene Q pure detection software (V2.3.1), were used for quantitative PCR (qPCR). The primers SZ35 (5’-AGGCTAAGCCGGGGTAAGTT-3’) and SZ36 (5’-GCCAAAAGCTTAAACTGCGG-3’) were used to quantify the nuclear DNA and the primers AA178 (5’-CATACCGAATAAACATCAGGGTAATCT-3’), AA179 (5’-ACGGGTGTTACACTATGATGAAGA-3’) were used to quantify the mtDNA levels. The cycle conditions were 95°C for 10□min, followed by 45 cycles of 95°C for 15□s and 58°C for 40□s. Primer efficiencies were determined by amplifying a series of exponential dilutions (1 to 10^−8^□ng) of cloned wild-type mtDNA (pSZ116) and *ges-1* promoter (pSZ57) fragments, confirming an equivalent efficiency of both sets of primers (91.2% for SZ35 and SZ36 and 90.6% for AA178 and AA179).

### Image acquisition and processing

Imaging was performed on live animals mounted on a 2% agarose pad on glass slides with 1 mM levamisole (Sigma). For quantification of purified mitochondria and mitochondrial membrane potential, we visualized fluorescence using a Zeiss Z2 imager microscope with a Zeiss Axiocam 506 mono camera and Zen2 (version 2.0.0.0) software. Mitochondria were counted with the aid of the Image J Grid plugin, and comparisons were made on the same immunoprecipitation volume for each sample. For 3D reconstruction, imaging/analysis was performed at the Queensland Brain Institute’s Advanced Microscopy Facility using a spinning-disk confocal system (Marianas; 3I, Inc.) consisting of an Aanio Observer Z1 (Carl Zeiss) equipped with a CSU-W1 spinning-disk head (Yokogawa Corporation of America), ORCA-Flash4.0 v2 sCMOS camera (Hamamatsu Photonics), and 20x 0.8 NA PlanApo and 40x 1.2 NA C-Apo objectives. Image acquisition was performed using SlideBook 6.0 (3I, Inc). 3D reconstruction and mitochondrial volume calculations were performed using 3D rendering in Imaris software (version 8.4.1, Bitplane). For all mitochondrial and mitophagy volume measurements, 100x magnification was used and representative images depict multiple images stitched together and straightened using image J.

### Mitochondrial membrane potential

Mitochondrial membrane potential was measured by observing mitochondrial accumulation of the fluorescent dye tetramethylrhodamine ethyl ester (TMRE; Sigma), as previously described [38, 39]. Briefly, 1-day old adult animals were transferred to growth plates seeded with 200 μl of M9 buffer containing 1 mM of TMRE. Animals were allowed to grow for 15 hours on the plates before being transferred to growth plates without TMRE for 5 hours to allow excess dye to be cleared from the intestinal cavity. Live animals were then prepared for fluorescence microscopy, as described above. To quantify the amount of dye accumulation in mitochondria from each tissue type, a region of interest was selected from images (40x magnification) containing mitochondria labelled with GFP fused to an MTS and driven under promoters that express in either body wall muscle (*myo-3*p::GFP^mt^) [40], or the PLM neuron (*mec-4*p::^MTS^GFP) [41]. Mean TMRE intensity was calculated for mitochondria of individual animals using Image J software.

### Statistics and reproducibility

The statistical analyses are described for each figure. Generally, an unpaired parametric two-tailed *t*-test was used to determine the significance of comparisons made between two data sets, unless otherwise noted in the figure legend. One-way analysis of variance (ANOVA) was performed for comparisons across multiple independent samples, using Tukey’s multiple comparisons correction. All experiments were reproduced at least 3 times with similar results. All data and biological reagents that support the conclusions of this manuscript are available from the corresponding author on request.

## Acknowledgements

We are grateful to M. Hilliard and R. Tweedale for comments on the manuscript and members of the Zuryn laboratory for discussions and comments. We also thank A. Gaudin for support with microscopy. Some strains were provided by the CGC, which is funded by the NIH Office of Research Infrastructure Programs (P40 OD010440). This work was supported by NHMRC Project Grants GNT1128381 (S.Z.) and GNT1162553 (S.Z.), ARC Discovery grants DP200101630 (S.Z.) and DP160103812 (J.G.), a Clem Jones Centre for Ageing Dementia Flagship grant (S.Z.), a Stafford Fox Senior Research Fellowship to S.Z., and a University of Queensland International Scholarship to C.Y.D.

## Author contributions

A.A. carried out most experiments. N.C., C.Y.D., S.Z. contributed some experiments. J.G. supervised N.C. and contributed to the interpretation of experiments. A.A., N.C., C.Y.D and S.Z. designed and interpreted experiments. A.A. and S.Z. wrote the paper. All authors approved the final manuscript.

## Supplemental Figure Legends

**Figure S1. ΔmtDNA heteroplasmy and genetic overview.** (**A**) Quantification of *uaDf5* heteroplasmy in whole animal extracts. Heteroplasmy was stable at around 50-60% for each of the strains used. Bar represents the mean value of three independent experiments ± s.e.m. (**B**) Schematic of *C. elegans* mtDNA showing the location of the 3.1kb *uadF5* deletion.

**Figure S2. Perturbing mitophagy increases mitochondrial volume in body wall muscle cells.** (**A**) Representative microimages of body wall muscle mitochondria. White scale bar, 50μm; red scale bar, 5μm. All images are stitched together from multiple high magnification (40x) z-stack images of live animals (see materials and methods). (**B**) Quantification of mitochondrial. Columns are means ± s.e.m. of individual animals (black dots). ** = *p*< 0.01. *P* values were determined using Student’s t-test.

**Figure S3. ΔmtDNA does not alter mitochondrial morphology or volume in body wall muscle cells and neurons.** Representative microimages of body wall muscle (**A**) and (**D**) motor neuron mitochondria. White scale bar, 50μm; red scale bar, 5μm. All images are stitched together from multiple high magnification (40x) z-stack images of live animals (see materials and methods). Quantification of mitochondrial volumes from (**B-C**) body wall muscle cells and (**E-F**) motor neurons. Columns are means ± s.e.m. of individual animals (black dots).

